# Rifampicin or capreomycin induced remodelling of the *Mycobacterium smegmatis* mycolic acid layer is mitigated in synergistic combinations with cationic antimicrobial peptides

**DOI:** 10.1101/269324

**Authors:** DeDe Kwun-Wai Man, Tokuwa Kanno, Giorgia Manzo, Brian D. Robertson, Jenny K.W. Lam, A. James Mason

## Abstract

The mycobacterial cell wall affords natural resistance to antibiotics. Antimicrobial peptides (AMPs) modify the surface properties of mycobacteria and can act synergistically with antibiotics from differing classes. Here we investigate the response of *Mycobacterium smegmatis* to the presence of rifampicin or capreomycin, either alone or in combination with two synthetic, cationic, α-helical AMPs; distinguished by the presence (D-LAK120-HP13) or absence (D-LAK120-A) of a kink-inducing proline. Using a combination of high-resolution magic angle spinning (HR-MAS) NMR of bacteria, diphenylhexatriene (DPH) fluorescence anisotropy and laurdan emission spectroscopy we show that *M. smegmatis* responds to challenge with rifampicin or capreomycin by substantially altering its metabolism and, in particular, by remodelling the cell envelope. In NMR spectra of bacteria, reductions in intensity for mycolic acid lipid −(CH_2_)-, -CH_3_, R_2_CH-COOH, R_2_CH-OH and also -CH_2_-(CH==CH)- and -CH=CH- resonances were observed following challenge with rifampicin and capreomycin, while the latter also caused an increase in trehalose. These changes are consistent with a reduction of trehalose dimycolate and increase of trehalose monomycolate and are associated with an increase in rigidity of the mycolic acid layer observed following challenge by capreomycin but not rifampicin. Challenge with D-LAK120-A or D-LAK120-HP13 induced no or modest changes respectively in these metabolites and did not induce a significant increase in rigidity of the mycolic acid layer. Further, the response to rifampicin or capreomycin was significantly reduced when these were combined respectively with D-LAK120-HP13 and D-LAK120-A, suggesting a possible mechanism for the synergy of these combinations. The remodelling of the mycomembrane in *M. smegmatis* is therefore identified as an important countermeasure deployed against rifampicin or capreomycin, but this can be mitigated, and rifampicin or capreomycin efficacy potentiated, by combining with AMPs.

## Introduction

*Mycobacterium spp*. are responsible for a variety of diseases including tuberculosis, leprosy, pulmonary disease, lymphadenitis, skin and disseminated diseases. Although tuberculosis incidence is falling globally at a rate of c 2% a year, it still carries the greatest worldwide disease burden with 10.4 million people falling ill with TB in 2016 and 1.7 million deaths.^1^ Infections due to non-tuberculous mycobacteria are also increasingly recognised.^2^ Although the reduced global incidence of tuberculosis is welcome, first line therapies for tuberculosis are increasingly failing with 600,000 new cases of tuberculosis resistant to the most effective first-line antibiotic, rifampicin. Of these, 490,000 cases were multi-drug resistant (MDR) and a substantial proportion of these are extensively drug-resistant (XDR). While 95% of tuberculosis deaths occur in low- and middle-income countries (LMICs), tuberculosis is also an increasing problem in the developed world where reactivation of latent tuberculosis infections is of particular concern.^3^ Mycobacteria are intrinsically resistant to many antibiotics that are effective against other bacteria and this is thought to be due to the superior protection offered by both mycobacterial outer (7-8 nm thick) and inner (6-7 nm) membranes, a layer of arabinogalactan-peptidoglycan (6-7 nm) and a periplasm (14-17 nm) containing lipomannan and lipoarabinomannan resulting in a cell envelope between 33 and 39 nm thick.^4,5^ The emergence of resistance to the limited number of existing therapies for tuberculosis has focussed attention on the particular problems associated with finding drugs capable of breaching the mycobacterial cell wall, since this may offer a means of either directly killing mycobacterial pathogens or of re-sensitizing them to existing first-line antibiotics.

In this regard, antimicrobial peptides are of considerable interest since their mechanisms of action are often associated with either direct damage to bacterial plasma membranes and/or penetration within the bacterial cytoplasm to access intracellular targets^6^, and these actions often result in rapid bacterial cell death - a property that may be desirable in reducing the incidence of resistance to AMPs. Antimicrobial peptides are therefore increasingly being evaluated as anti-tuberculosis agents.^7,8^

**Figure 1.**
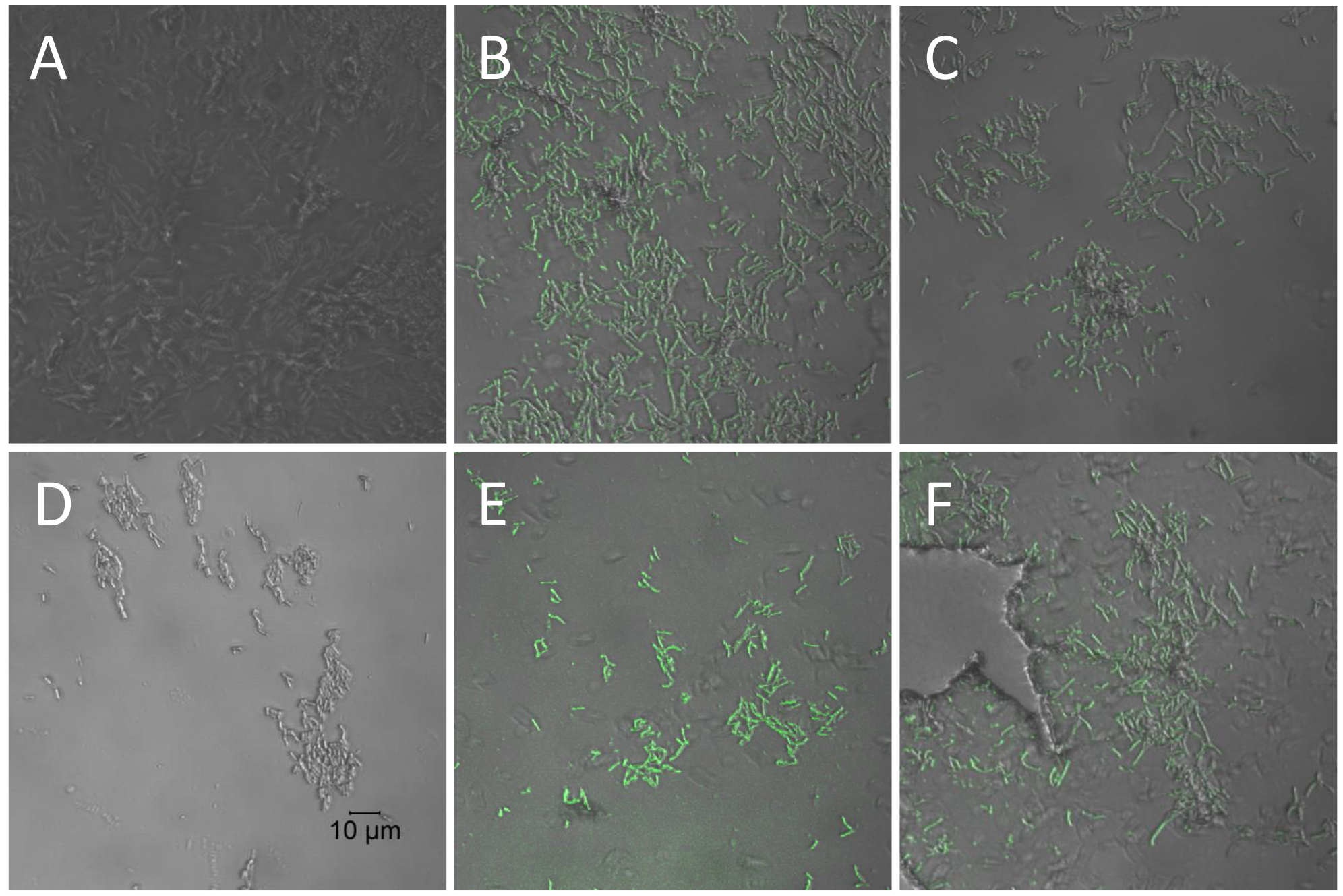
AMPs render *Mycobacterium smegmatis* permeable to fluorescein isothiocyanate (FITC) - dextran. Confocal microscope images of 1.1 × 10^8^ CFU/ml *M. smegmatis mc*^2^ 155 either unchallenged (A) or challenged with rifampicin (D), ½ MIC (B/C) or 2 x MIC (E/F)), D-LAK120-A (B/E) or D-LAK120-HP13 (C/F). In contrast to rifampicin, both peptides facilitate absorption or adsorption of the 150 kDa fluorescently labelled dextran.

Indeed, not only have antimicrobial peptides been evaluated in isolation, combinations of cationic α-helical peptides with first line anti-mycobacterial agents have already been shown to be effective.^9^ All D-amino acid isomers of these peptides displayed improved stability and enhanced mycobacterial selectivity.^10^

Our own research has focused on a series of highly cationic antimicrobial peptides, comprising D-amino acids, rationally designed to adopt α-helix conformations within biological membranes.^11^ These AMPs have a detergent like ability enabling disruption of colonies of *Mycobacterium tuberculosis* H37Ra^11^, are able to inhibit growth of MDR and XDR strains of *M. tuberculosis* when cultured in THP-1 macrophage and potentiate the activity of first-line antibiotic isoniazid *in vitro*.^12^ The D-LAK peptides were designed to be cationic and amphipathic, with the angle subtended by the positively charged lysine residues when the peptide adopts an idealised α-helix conformation, modified to enhance disruption of anionic model membranes.^11^ Furthermore, the role of conformational flexibility was investigated and proline residues were introduced to disrupt the α-helix. The positioning of this proline kink was shown to be important and could enhance activity against Gram-negative bacteria while also mitigating haemolysis.^11^ Conformational flexibility is a key property of potent antimicrobial peptides, such as pleurocidin, which is associated with greater penetration of bacterial membranes and the ability to reach intracellular targets.^12,13^ A transcriptomic and NMR metabolomic approach to understanding the mechanism of action of D-LAK120-HP13 supports an ability to penetrate within Gram-negative bacteria.^13^ Interestingly however the proline containing D-LAK peptides did not always outperform their proline free analogues when tested against *Mycobacterium tuberculosis* H37Ra highlighting that the mechanism of action against mycobacteria is likely to be distinct from that which is effective against Gram-negative bacteria. Consequently, our understanding of the structural features that promote anti-mycobacteria activity is poorly developed.

To address this gap in our understanding we have investigated in more detail how *Mycobacterium smegmatis* responds to challenge with proline free (D-LAK120-A) and proline containing (D-LAK120-HP13) D-LAK peptides. We examined the activity of, and bacterial response to, combinations of each peptide with rifampicin or capreomycin to investigate the mechanism of the observed, albeit modest, synergy. Testing the hypothesis that the membrane activity of the D-LAK peptides is key to both their anti-mycobacterial activity and their ability to potentiate the activity of rifampicin and capreomycin, we incorporate high resolution magic angle spinning (HR-MAS) NMR metabolomics with fluorescence spectroscopy of *M. smegmatis* membrane-incorporated trans-1,6-diphe-nyl-1,3,5-hexatriene (DPH) and laurdan dyes which are sensitive to changes in membrane fluidity and order, respectively. These techniques are able to distinguish between the two D-LAK peptides and reveal that, when challenged with sub-inhibitory concentrations of rifampicin or capreomycin, *M. smegmatis* remodels its membrane. These observations suggest why and how antimicrobial peptides may act and may be modified to improve their ability to potentiate existing anti-mycobacterial treatments.

## Materials and Methods

*Mycobacterium smegmatis* mc^2^ 155 was grown in Middlebrook 7H9 broth (Difco, Detriot, Michigan, USA, Cat.# 271310) supplemented with 10% (v/v) oleic acid-albumin-dextrose-catalase (OADC) enrichment (Becton Dickinson, Franklin Lakes, New Jersey, UsA, Cat.# 212240) and 0.5% (v/v) glycerol (Sigma Aldrich, Cat.%G-5516) in 50 ml Falcon tubes at 37°C without shaking. 0.025% (v/v) Tyloxapol surfactant (Sigma Aldrich, Cat.# T8761) was added as one of the treatments. Rifampicin (RIF; R3501) and capreomycin sulfate (CAP) from *Streptomyces capreolus* (C4142) were from Sigma Aldrich. D-AMPs D-LAK120-A *(KKLALALAKKWLALAKKLALALAKK*-NH_2_) and D-LAK120-HP13 (*KKALAHALKKWLPALKKLAHALAKK*-NH_2_) were supplied by China peptides Co. Ltd. (Shanghai, China). The purity of synthesized peptides was above 80% and the peptides were used as supplied. Both peptides were amidated at the C-terminus.

*Minimum inhibitory concentration (MIC) and Fractional inhibitory concentration (FIC)* MIC of the anti-TB agents (rifampicin, DAMPs and capreomycin) against *M. smegmatis* mc^2^ 155 was determined using broth micro-dilution assay in 96-well plates. Two-fold serial dilutions were made in Middlebrook 7H9 broth supplemented with oleic albumin dextrose catalase (OADC) and glycerol in duplicate. An inoculum at an OD_600_ of 0.02 was prepared by diluting mid-log cultures and 100 μl was added in each well (~1 x 10^5^ cfu). D-AMPs were prepared at 128 μM to 1 μM in Middlebrook broth. RIF was serially diluted from 100 μg/ml to 0.195 μg/ml. Capreomycin was were prepared at 5 μg/ml to 0.004 μg/ml. 100 μl of mycobacterial suspension and anti-TB agent were added in each well to obtain the inoculum concentration of 5 × 10^4^ CFU/ml. Growth control without drug and sterile medium were prepared in each assay. The plates were then incubated at 37°C for 24h before adding 30 μl of 0.02% (w/v) resazurin dye and the colour change was evaluated after incubation for another 24h. Resazurin assay is a colorimetric assay in which viable cells would reduce blue resazurin (oxidized state) into fluorescent pink resorufin (reduced state). MIC of each anti-TB agent was determined as the lowest concentration that prevented growth, and therefore remained blue.

Fractional inhibitory concentration (FIC) of the combination treatment of rifampicin or capreomycin and D-AMPs was determined using a checkerboard assay.^15^ Synergistic interactions between anti-TB agents in combination treatment was denoted by the Fractional inhibitory concentration (FIC) index. FIC values of each combination were calculated from the following formula:

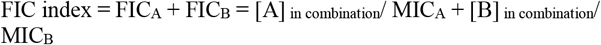

Where [A] is the lowest inhibitory concentration of D-AMPs in the presence of RIF or CAP, MICA is the MIC of D-AMPs treated as a single agent and FICA is the FIC of D-LAK peptides; [B], MICB and FICB are the corresponding values for RIF or CAP. Resembling a coordinate plane, the 96-well plate was set into x- and y-axis which were designated for the addition of D-AMPs and RIF or CAP respectively. Two-fold serial dilutions of each anti-TB agent [x-axis: D-AMPs (4 μM to 0 μM); y-axis: RIF (100 μg/ml to 0 μg/ml); CAP (5 μg/ml to 0 μg/ml)] was added into the designated wells to give combination treatments as follows: D-LAK120-A; RIF, D-LAK120-A; CAP, D-LAK120-HP13; RIF and D-LAK120-HP13;CAP. Mycobacterial suspension was prepared as described above and 1 ×x 10^5^ CFU/ml was seeded in each well. After 24h, the resazurin assay was performed and FIC was determined as the lowest combination concentrations that remained blue.

*Confocal microscopy. M. smegmatis* mc^2^ 155 (1.1 × 10^8^ CFU/ml) mid-log phase (O.D. =;0.6) was treated with 250 μg/ml 150 kDa FITC-labelled dextran (Sigma Aldrich, Cat. #46946) and D-AMPs (D-LAK 120 A or D-LAK 120 HP13) or rifampicin at the respective MICs for 20 min. Following treatment, the bacterial cells were centrifuged at 7500 rpm for 10 min and washed three times with PBS. Cells were then fixed in 2% formaldehyde at R.T for 20 min. After fixation, cells were washed twice and re-suspended in PBS. 20 μl of sample were then allowed to air-dry on microscope slides overnight. Samples were then imaged with a 65 x oil-immersion objective lens using a Zeiss LSM-510 inverted confocal microscope (Carl Zeiss Inc.) FITC was excited with a 488 nm laser and detected with a 505 nm long-pass filter.

*Transmission electron microscopy* (TEM). *M. smegmatis* mc^2^ 155 (1 × 10^7^ CFU/ml) at mid-log phase (O.D. = 0.6) was treated with D-LAK120-A and D-LAK120-HP13 at their MICs respectively for 5 and 30 min. Samples were washed three times by PBS and cell pellets were collected by centrifugation. Cell pellets were then fixed in 2.5% glutaraldehyde till further processing. Samples were rinsed thrice with piperazine-N,N’-bis(ethanesulfonic acid); 1,4-piperazinediethanesulfonic acid (PIPES buffer) (Sigma Aldrich Cat.# P6757) followed by second fixation in 1% osmium tetroxide (OsO4) (Sigma Aldrich Cat.# 75632) for 1 h at R.T. 1^st^ embedding into agar was done to reduce sample loss. Pellets were then dehydrated using ethanol (EtOH) (50, 70, 90% for 10 min each and 3 × 100% for 20 min). After dehydration, samples were infiltrated using 1:1 epoxy resin/ propylene oxide mixture overnight at 37 °C. The next day, pellets were infiltrated with fresh epoxy resin for 1h at 37 °C followed by polymerization at 60 °C overnight in plastic moulds. Samples were viewed using Philips CM100 TEM equipped with a TENGRA 2.3K X 2.3K camera.

*Anti-TB agent challenge assays*. To evaluate the mechanism of action of anti-TB agents on bacterial growth, a ¾ MIC challenge assay was conducted to observe the growth response of*M. smegmatis* mc^2^ 155. 1% inoculum in the presence of ¾ MIC of each anti-TB agent, alone or in combination, was cultured in 10 ml Middlebrook broth medium at 37°C. After 5 days, the bacteria were processed and subjected to trans-1,6-diphenyl-1,3,5-hexatriene (DPH) fluorescence assay, laurdan fluorescence emission and nuclear magnetic resonance (NMR) metabolomics analyses.

*Trans-1,6-diphenyl-1,3,5-hexatriene (DPH) fluorescence assay. M. smegmatis* mc^2^ 155 bacterial suspension was harvested and fixed by 0.25% formaldehyde for 1 h at room temperature. After fixing, the bacterial cells were centrifuged at 5000 rpm, 4 °C for 8 min then washed by 5 ml phosphate buffered saline (PBS) once. Bacteria were re-suspended in PBS and stained by 2.5 μM DPH fluorescence probe. All tubes were wrapped with foil to prevent light degradation of the dye and incubated at 37 °C for 30 min. Samples were added into the cuvette and excited at 358 nm using a Varian Cary Eclipse Fluorescence Spectrophotometer. Signals were collected by single wavelength scanning at emission wavelength 430 nm. The spectral bandwidth of the emission monochromator and the measurement of emission spectra are set at 10 nm. Steady-state fluorescence anisotropy, r, was calculated as

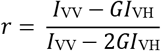

where I_VV_ and I_VH_ are the parallel and perpendicular polarized fluorescence intensities measured with the vertically polarized excitation light, I_HV_ and I_HH_ are the same fluorescence intensities measured with the excitation light horizontally polarized, and G is the monochromator grating correction factor given by G = I_HV_/I_HH_.

*Laurdan fluorescence assay*. Laurdan (6-dodecanolyl—N,N-dimethyl-2-napthtylamine) *M. smegmatis* mc^2^ 155 bacterial suspension was harvested and fixed by 0.25% formaldehyde for 1 h at room temperature. The bacterial pellet was then fixed, washed and resuspended as stated previously. The suspension was then stained by 2.5 μM Laurdan fluorescence probe. All tubes were wrapped with foil to prevent light degradation of the dye and incubated at 37 °C for 1 h. Samples were added into the cuvette and excited at 350 nm using varian Cary Eclipse Fluorescence Spectrophotometer. Signals were collected at emission wavelength range of 400 to 600 nm (no. of scans = 20). General polarization (GP) value corresponding to the fluidity of membrane was calculation as below where I_440_ and I_490_ are the emission intensities at corresponding wavelengths:

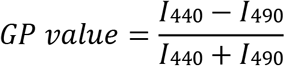

*NMR metabolomics. M. smegmatis* mc^2^ 155 bacterial suspension were pelleted by centrifugation at 5000 rpm, 4 °C for 8 min. Supernatant was collected and filtered through 0.22 μm membrane filters to remove remaining bacterial cells. Bacterial pellet was washed twice and re-suspended in PBS. All supernatant samples were frozen at −80 °C and pellets were snap frozen in liquid nitrogen before freeze drying using an Alpha 1-2 LD plus freeze dryer (Martin Christ, Germany). After lyophilization, the supernatant was rehydrated by 10% D_2_O containing 2,2,3,3-D_4_-3-(Trimethylsilyl) propionic acid sodium salt (TMSP-2,2,3,3-D_4_) to provide a deuterium lock signal with reference signal. Samples were then transferred to NMR tubes. ^1^H NMR spectra were recorded on a Bruker Avance II 700 NMR spectrometer (Bruker BioSpin, Coventry, United Kingdom) equipped with a 5-mm helium-cooled quadruple resonance cryoprobe with sample isolates tested (9 replicates) and kept at 4 °C. 1D spectra were recorded under automation at 298K using a Carr-Purcell-Meiboom-Gill pre-saturation (cpmgpr1) pulse sequence. Spectra were acquired with 64 transients, a spectrum length of 20.1 ppm and 65 k data points. For the freeze dried bacterial pellets, 40 μl of D_2_O was used for rehydration. The samples were introduced in the HRMAS probe on a 600 MHz Bruker Avance Ill spectrometer, keeping the temperature at 310 K. Spinning speed was 5 kHz. ^1^H NMR spectra were collected with a Carr-Purcell-Meiboom-Gill pre-saturation (zgpr) pulse sequence, a spectrum width of 12.01 ppm and 16384 data points was used. A total of 128 scans were used to obtain each of the NMR spectra. Free induction decay was multiplied by an exponential function with line broadening of 0.3 Hz. To aid the assignment of metabolite resonance, correlation spectroscopy (COSY) (cosygpprqf) spectra were acquired for a subset of samples. All peak positions were Fourier transformed and measured relative to the trimethylsilylpropanoic (TSP) acid-D_4_ reference peak set to 0.0 ppm. Phase correction of spectra was performed manually, and automatic baseline correction was applied.

*Data analysis and statistics*. All microbial data presented in this study were statistically analysed by GraphPad Prism (version 5.01 for Windows, GraphPad Software, La Jolla California USA). Oneway ANOVA test was performed, followed by Dunnett’s Multiple Comparison Test with a value of p ≤ 0.05 to establish statistical significance. Sigmodal dose-response (variable slope) was used for fitting of growth inhibition calculations. All experiments were performed for at least 3 times.

For metabolomics data, peak assignment was performed by comparing with chemical shift values and multiplicities from J-resolved NMR spectra to values from the BMRB,^16^ COSY spectra and Chenomix NMR suite software (Chenomx Inc., Edmonton, Canada) with NMR metabolite templates. Multivariate data analysis was based on previous published work.^17^ Principal component analysis (PCA) and orthogonal partial least squares discriminant analysis (OPLS-DA) were done using MVAPACK^18^ and a software developed in our previous study.^17^ The software involves the use of Python programming language with NumPy and SciPy for calculations and visualization is performed by matplotlib. To minimize the noise content, regions beyond 10.5 ppm, below 0 ppm and water peak were excluded. These spectra were subjected to probabilistic quotient normalization (PQN) and autoscaled.^19^ Icoshift algorithm was applied for further alignment and optimized bucketing algorithm with a 0.005 ppm bin size was employed for bucketing. Cross-validation was performed where 85% of the samples were used as a training set and the remaining 15% as a test set, ensuring that the number of samples in the test set was proportional to the total number of samples from each class, while at least one sample from each class was present in the test set. The F1-score and leave-one-out cross-validation was carried out on the samples in the training set in order to choose the number of components for the model, with the additional constraint to use a maximum of 5 components. This double cross-validation was repeated 1000 times with randomly chosen samples in the training and test set to prevent bias due to the choice of training or test set. This produces 7×1000 models (in the supplementary information, each of these models leads to a point on the scores plot, but loadings and weights are presented as averages over all these models). Reference Q^2^ value was generated after the known or random class assignment process. It provides a measure of goodness of fit after cross validation and generally referred to be ‘good’ above 0.5.^20,21^ The Q^2^ value quoted is the mean of all models and was compared between the genuine and permutated class assignments in each case. Backscaled loadings plots were used to identify metabolites with high variance and weight compared with the untreated control spectra after PQN normalization. Heatmaps with Euclidian distance-hierarchical cluster (HCL) analysis were generated by MultiExperiment Viewer (MeV) software.

*Volcano plots and univariate analysis of metabolomics data*. For volcano plots, significant results from PLS or OPLS-DA analysis justified univariate analysis of peaks associated with predictive value in the multivariate models. Therefore, the integrals of assigned NMR peaks were analysed using non-parametric univariate methods. Volcano plots were compared the fold change in metabolite values between two conditions. Fold change was calculated as the ratio between the treatment condition and the control (Treatment/Control). Mann-Whitney U test were used to compare means and associated p-values were False Discovery rate adjusted using the Benjamini-Hochberg method (α = 0.05). Volcano plots were generated using custom scripts in Python with Numpy, Pandas, Matplotlib and Seaborn packages.

*PLS regression* - PLS Regression was performed in Python using the PLSRegression function in the scikit-learn software package. Initial model assessment determined that only one component was necessary for best model performance. Increased number of components led to a decrease in Q^2^. Monte Carlo cross-validation of models was performed by randomly splitting data into 70/30 training/test set splits. Model generation and assessment was repeated 1000 times to avoid bias by sample separation and R^2^ and Q^2^ values were calculated to assess model performance. The R^2^ metric demonstrates how well the model describes the training data set while Q^2^ is a metric of how well the model predicts the test set.

**Figure 2.**
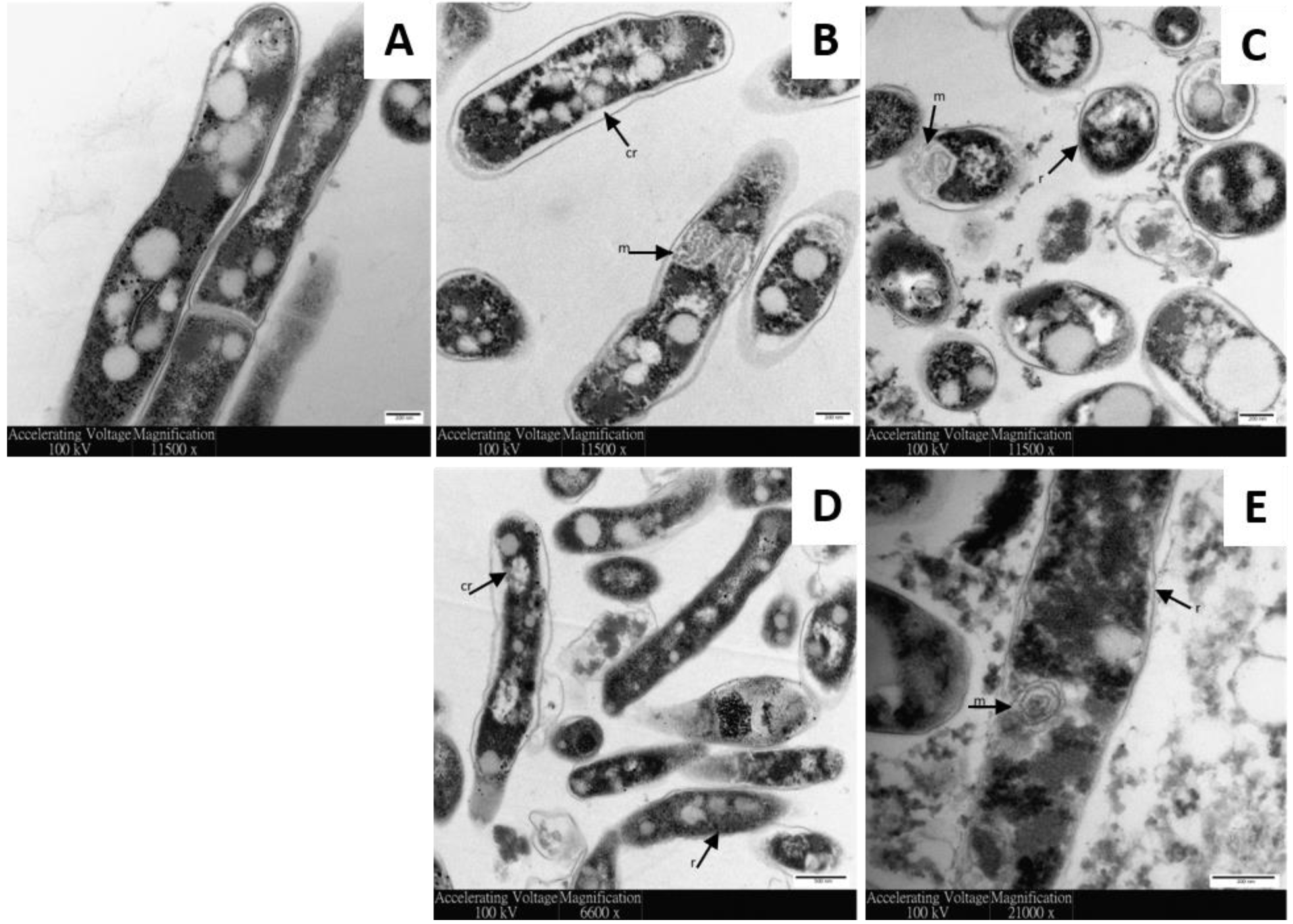
AMPs disrupt membrane surface of *M. smegmatis*. Transmission electron micrographs of 1.1 × 108 CFU/ml M. smegmatis mc2 155 either unchallenged (A) or challenged with D-LAK120-A (B/C) or D-LAK120-HP13 (D/E) for 5 (B/D) or 30 (C/D) mins. r ‒ membrane ruffling, cr ‒ cytoplasmic retraction, m ‒ mesosoma.

## Results

*Action of D-LAK peptides on M. smegmatis*. The action of the two D-LAK AMPs on *M. smegmatis* was first investigated to confirm that, as suspected from their activity against *M. tuberculosis*, the peptides acted on the mycobacterial cell wall. Confocal microscopic investigation of *M. smegmatis* mc^2^ 155 challenged with sub- or supra-MIC concentrations of D-LAK120-A, D-LAK120-HP13 or rifampicin indicate that both peptides, but not rifampicin, cause the bacteria to stain positive with the FITC labelled 150 kDa dextran (Fig. 1). FITC-dextran has been used in a number of studies with vesicles to probe the membrane permeabilization ability of diverse antimicrobial peptides, and a series of labelled dextran conjugates are available to measure the size of pores that may be formed by antimicrobial peptides.^22–25^ In the case of melittin, dextran of 4 kDa was able to readily escape from loaded vesicles but much less of the 50 kDa analogue was released. This indicated a pore with a defined diameter of 25-30 Å is formed.^22^ In contrast, other peptides facilitate escape of dextrans across the full range of sizes tested but complete leakage is never achieved.^24,25^ These results have been interpreted as evidence for rather large but transient lesions.^24^ At 150 kDa the dextran used in the present study is substantially larger than those used in previous studies of escape from lipid vesicles and it is not possible from images with this resolution to determine whether the labelled dextran has penetrated within the bacteria. It is may be that the same functionality that causes D-LAK peptides to prevent aggregation of*M. tuberculosis* colonies also enables the dextran to adsorb to the surface of *M. smegmatis*.

Transmission electron microscopy was exploited to visualize the effect of D-LAK peptides on the ultrastructural changes of *M. smegmatis* (Fig. 2). Unchallenged cells have characteristic lipid inclusions, a regular rod-shaped cell body with well-defined intact cell membranes and homogeneous cytoplasm (Fig. 2A). The cell envelope was distinctly visible with cytosol bound by the plasma membrane, surrounded by an internal and then an outer electron dense layer.^26,27^ The cell content was slightly pulled away in some control cells which is attributed to sample preparation process. Membrane ruffling was observed after treating *M. smegmatis* mc^2^ 155 at 1 × MIC with D-LAK120-A (Fig. 2B) or D-LAK-120-HP13 (Fig. 2D) for 5 mins. Cytoplasmic retraction was observed as the inner cell membrane detaches from the outer cell wall. This phenomenon was reported when *Escherichia coli* and *Bacillus subtilis* were exposed to antimicrobial peptides.^28^ Similar findings have been observed for exposure of *Mycobacteria tuberculosis* (Erdman strain) to antimicrobial peptides; granulysin demonstrated inner cell membrane detachment as observed in osmotic lysis.^29^ The reduction in membrane uniformity suggests a cell penetrating ability for the D-LAK peptides. Peptide treatment also induced notable intracellular changes, with clumped cytosol and increased amount of mesosoma formation. Mesosome structures have been observed previously in bacteria challenged by antimicrobial peptides.^30,31^ Formation of mesosomes was suggested as a repair mechanism by bacteria to combat cell lysis.^32^ Prolonged treatment (30 min) by D-LAK peptides resulted in large scale of cell lysis and presence of cell debris (Fig. 2C/E). This indicates that the fate of bacteria challenged with inhibitory concentrations of D-LAK peptides is disturbed cell membranes and cell wall disintegration, membrane rupture and release of intracellular content.

**Table 1.** Minimal inhibitory concentrations of D-LAK peptides, rifampicin and capreomycin against *M. smegmatis* mc^2^ 155 in the presence or absence of tyloxapol.

|  | MIC<sub>50</sub> (μM) |  |
| --- | --- | --- |
|  | Tyloxapol | No tyloxapol |
| Rifampicin | 19.9 ± 3.04 | 38.2 ± 5.83 |
| Capreomycin | 1.72 ± 0.43 | 1.34 ± 0.25 |
| D-LAK120-A | 0.82 ± 0.07 | 1.68 ± 0.43 |
| D-LAK120-HP13 | 0.63 ± 0.03 | 1.80 ± 0.39 |

*Activity of D-LAK peptides against* M. smegmatis *alone or in combination with rifampicin or capreomycin* - All four antibiotics were tested in isolation against *M. smegmatis* mc^2^ 155 both in the presence and absence of tyloxapol; a non-ionic liquid polymer which is often included when growing *Mycobacteria* in planktonic suspension to prevent clumping (Table 1). Notably both peptides and rifampicin were more potent when tyloxapol was present in the culture media but the activity of capreomycin was unchanged.

**Table 2.** Fractional inhibitory concentrations of D-LAK peptides in combination with either rifampicin or capreomycin against *M. smegmatis* mc^2^ 155 in the presence or absence of tyloxapol.

|  | FIC |  |
| --- | --- | --- |
|  | Tyloxapol | No tyloxapol |
| Rifampicin/D-LAK120-A | 1.58 | 2.50 |
| Rifampicin/D-LAK120-HP13 | 1.64 | 0.81 |
| Capreomycin/D-LAK120-A | 0.45 | 0.71 |
| Capreomycin/D-LAK120-HP13 | 0.53 | 0.97 |

Combinations of rifampicin or capreomycin were then tested with each of the D-LAK peptides (Table 2). In the absence of tyloxapol, the combinations of capreomycin and the D-LAK peptides are modestly synergistic. Effectively inhibition is achieved by D-LAK120-A and capreomycin when used in combination with approximately a third of each agent required to achieve the same effect as when each agent is used alone. The effect is more marked when the combination is used in the presence of tyloxapol with an FIC <0.5 which is widely accepted as a threshold for synergism. In these conditions approximately only one tenth of the amount of D-LAK120-A required for the same effect. A similar effect is seen when capreomycin is used in combination with D-LAK120-HP13 but the synergistic effect is more modest in both the presence and absence of tyloxapol. Rifampicin demonstrates no synergy with D-LAK120A and these antibiotics may even be antagonistic when no tyloxapol is added to the growth media. In contrast, though modest, D-LAK120-HP13 may assist the activity of rifampicin and a little less than half as much rifampicin is required to achieve the same inhibitory effect as when rifampicin is used alone. Analogous experiments were performed with isoniazid but no synergism was detected with either peptide and hence these combinations were not studied further in the present work.

*DPH anisotropy and laurdan fluorescence indicate substantial changes in membrane physical properties when* M. smegmatis *is cultured in the presence of capreomycin or tyloxapol* - DPH fluorescence polarization or fluorescence anisotropy is commonly used to determine changes in bacterial cytoplasmic membrane fluidity under environmental stress.^33^

**Figure 3.**
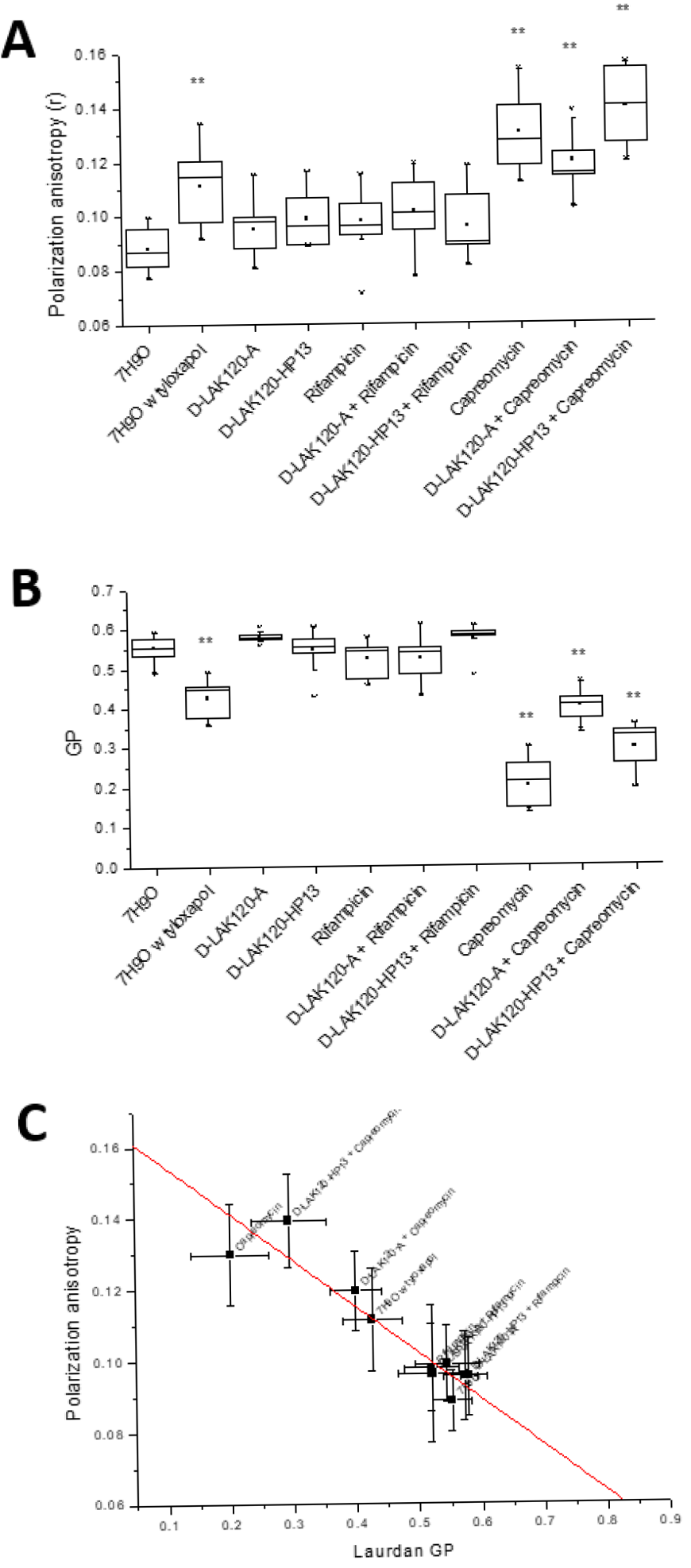
Fluorescence spectroscopic perspective of *M. smegmatis* mc^2^ 155 response to challenge with antibiotics. DPH fluorescence anisotropy (**A**), laurdan GP fluorescence (**B**) and correlation (R^2^ = 0.856) of DPH and laurdan (**C**). ** indicates *p* <0.05 with respect to 7H9O.

As a polyene hydrophobic dye DPH localises to the hydrophobic core of lipid bilayers. It aligns parallel to the acyl chains of lipids and hence its ability to re-orientate is dependent on the packing of its lipid neighbours. A high degree of orientation, and hence fluorescence anisotropy, results from a more rigid lipid bilayer. Significant increases (p < 0.05) in DPH fluorescence polarisation anisotropy (r) were observed for *M. smegmatis* mc^2^ 155 challenged during growth with ¾ MIC or FIC capreomycin or its combinations with D-LAK120A or D-LAK120-HP13 (Fig. 3A) or when grown in the presence of 0.025% tyloxapol; the latter producing a more modest increase. Small increases in fluorescence anisotropy were observed when bacteria were challenged with the D-LAK peptides or with rifampicin, either alone or in combination with the D-LAK peptides, but these were not significant when compared with unchallenged bacteria. Only challenge with capreomycin alone or in combination with either D-LAK peptide, causes a substantial increase in membrane rigidity in *M. smegmatis*.

Laurdan is a solvatochromic fluorescent dye which inserts into lipid bilayers with the fatty acid chain embedded in the hydrophobic core of the bilayer and the N,N-dimethyl-2-napththylamine in the more polar interfacial region. The dipole moment between the dimethyl-amino and carbonyl moieties is sensitive to local changes in polarity which will shift the emission maximum. Calculating the difference in emission at 440 and 490 nm generates a generalised polarization (GP) value from which membrane order can be inferred. As a lipid bilayer becomes disordered more water can penetrate the interfacial region, which becomes more polar, and the GP value decreases.^34^

The GP value is therefore a measure of disorder in the lipid bilayer interfacial region although other factors may influence the polarity of the environment in which the fluorophore is located and some caution is advisable when interpreting laurdan GP in biological systems. Nevertheless, laurdan generalised polarization has been used in e.g. *Bacillus subtilis* to detect the effect of challenge with a synthetic cyclic hexapeptide cWFW^35^ and the organisation of the bacterial membrane by flotillins^36^ and MreB^37^. Significant changes (*p* < 0.05) in laurdan GP were observed for *M. smegmatis* mc^2^ 155 challenged during growth with ¾ MIC or FIC capreomycin or its combinations with D-LAK120A or D-LAK120-HP13 (Fig. 3B). Again, a change in membrane properties was detected for bacteria incubated with tyloxapol but a noticeable reduction in GP following challenge with rifampicin is non-significant. The change in GP for bacteria challenged with combinations of capreomycin is of a lower magnitude than that observed when the bacteria are challenged with capreomycin alone but the differences between these three conditions are non-significant. Notably the GP values in each of conditions were reduced relative to the unchallenged bacteria, which implies the local environment of the laurdan probe is becoming more polar, something that would normally be associated with an increase in disorder in the interfacial region. Since the increase in DPH anisotropy for these conditions is associated with an increase in membrane rigidity, a less ordered but more rigid membrane is perhaps surprising. The strong correlation between DPH and laurdan data (Fig. 3C) does however indicate the two techniques are reporting on the same event. The DPH dye would be expected to reside deep within the hydrophobic core of the membrane while the laurdan probe would be much closer to the aqueous surface. Consequently, it is possible that the result of*M. smegmatis* responding to the presence of capreomycin is to modify the hydrophobic core of the membrane, reducing fluidity, while the interfacial region becomes more disordered. While the precise locations of both dyes in the *Mycobacteria* cell membranes are yet to be determined it can be concluded that the physical properties of the bacterial envelope are substantially altered in response to challenge with capreomycin but not with either rifampicin or the D-LAK peptide.

*HR-MAS^1^H NMR metabolomics reveals substantial changes in* M. smegmatis *membrane components* - Cross-validated OPLS-DA was used to identify significant changes in ^1^H HR-MAS spectra obtained for *M. smegmatis* mc^2^ 155 when challenged with ¾ MIC of individual anti-*mycobacteria* drugs, ¾ FIC of their combinations or 0.025% tyloxapol. The technique could determine significant differences, as determined by Q^2^ (Table 3), for each model in which loadings are compared in a hierarchical clustered heatmap (Fig. 4A). Volcano plots for individual comparisons allow the identification of the magnitude and significance of changes in individual metabolites (Fig. 4B-F) while changes in key metabolites are compared across treatments to show the impact of each peptide, rifampicin, capreomycin and their combinations (Fig. 5).

**Table 3.** Comparison of test and permutated Q^2^ scores for crossvalidated OPLS-DA models comparing ^1^H HR-MAS spectra of unchallenged *M. tuberculosis* mc^2^ 155 with those obtained for bacteria in the indicated conditions.

|  | Q<sup>2</sup> | Q<sup>2</sup><br>(permuted) |
| --- | --- | --- |
| Tyloxapol (0.025%) | 0.866 | -0.377 |
| D-LAK120-A | 0.769 | -0.423 |
| D-LAK120-HP13 | 0.729 | -0.360 |
| Rifampicin | 0.581 | -0.426 |
| Rifampicin/D-LAK120-A | 0.430 | -0.429 |
| Rifampicin/D-LAK120-HP13 | 0.763 | -0.410 |
| Capreomycin | 0.881 | -0.347 |
| Capreomycin/D-LAK120-A | 0.775 | -0.367 |
| Capreomycin/D-LAK120-HP13 | 0.823 | -0.344 |

Culturing *M. smegmatis* mc^2^ 155 in the presence of 0.025% tyloxapol causes significant and substantial changes in several metabolites and components of the mycobacterial membrane (Fig. 4B). An increase in the amount of glutamate in the cells is the greatest change while there are highly significant reductions in lipid resonances associated with mycolic acid including the saturated alkyl chains ‒(CH_2_)_n_- and both R_2_CH-COOH and R_2_CH-OH. Notably, resonances associated with unsaturated alkyl groups ‒CH=CH- or ‒CH_2_-CH=CH- are unaffected (Fig. 4B, 5A-D). Mycobacterial mycolic acids comprise a long branch mero chain of 40-60 carbons and a short α branch of typically 24 carbons. The shorter α chain is completely saturated. Mero-mycolic acids in other *Mycobacteria* contain cyclopropanated mycolic acids but these are uncommon in *M. smegmatis* where the major mycolic acids are a homologous, α’series containing just a *cis*-alkene in the mero-chain.^38–42^

Since the fold changes in all four lipid resonances are similar, this is consistent with a reduction in the number of saturated α chains, with mero-mycolic acids, containing unsaturated hydrocarbons, unaffected. Many of these differences are also observed when *M. smegmatis* mc^2^ 155 is challenged with either rifampicin or capreomycin (Fig. 4E/F).

**Figure 4.**
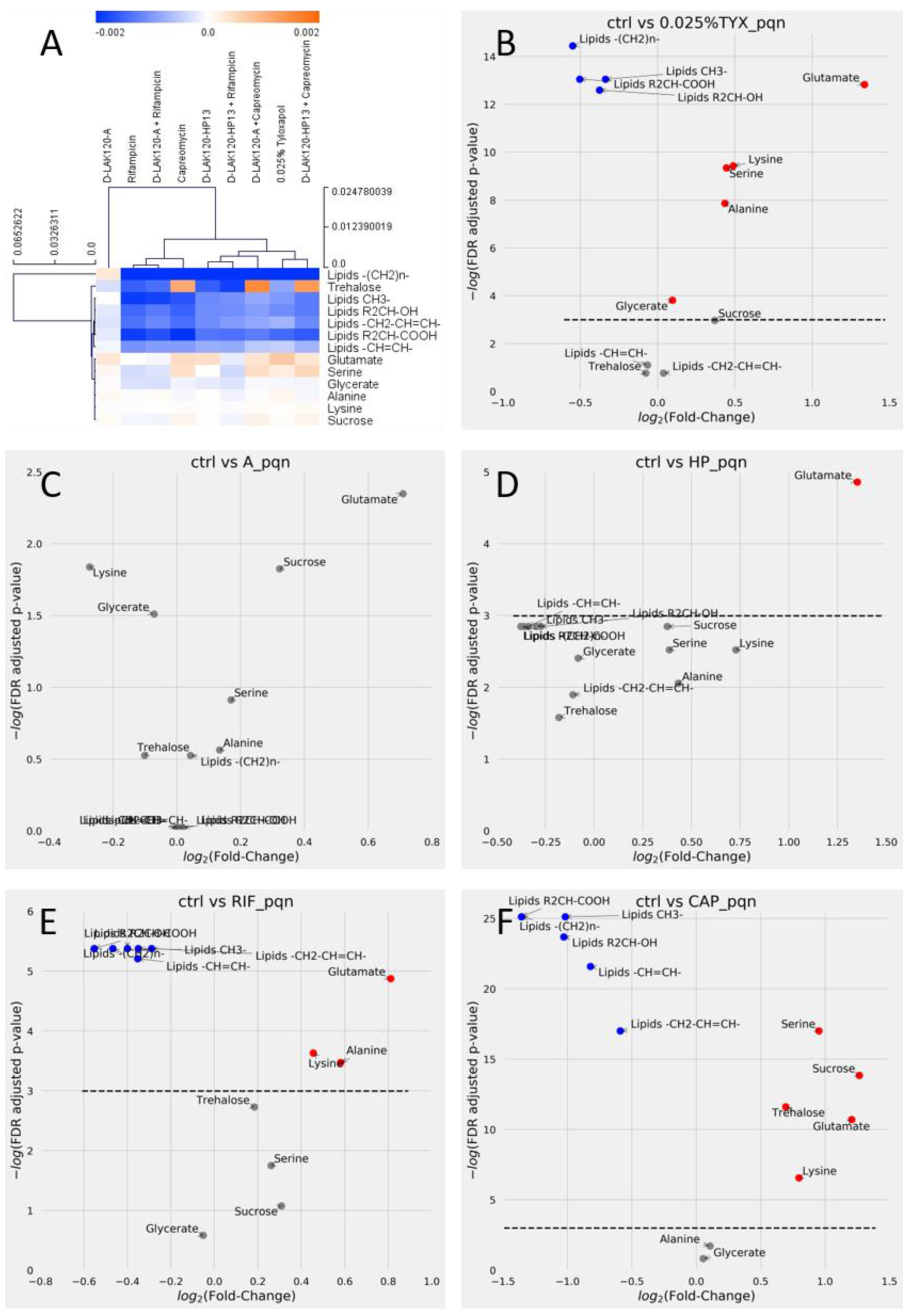
Capreomycin and rifampicin induce substantial changes in mycolic acid lipid components in *M. smegmatis*. Hierarchical clustered heatmap comparing loadings obtained from cross-validated OPLS-DA of ^1^H HR-MAS NMR spectra of *M. smegmatis* mc^2^ 155 grown in the indicated conditions (A). volcano plots are shown for individual comparisons of unchallenged bacteria and those challenged with 0.025% tyloxapol (B), D-LAK120-A (C), D-LAK120-HP13 (D), rifampicin (E) or capreomycin (F). Volcano plots are of PQN normalized data and allow comparison of fold changes and significance of each metabolite; blue - significant reductions, red - significant increases and grey - non-significant changes in the indicated metabolites.

**Figure 5.**
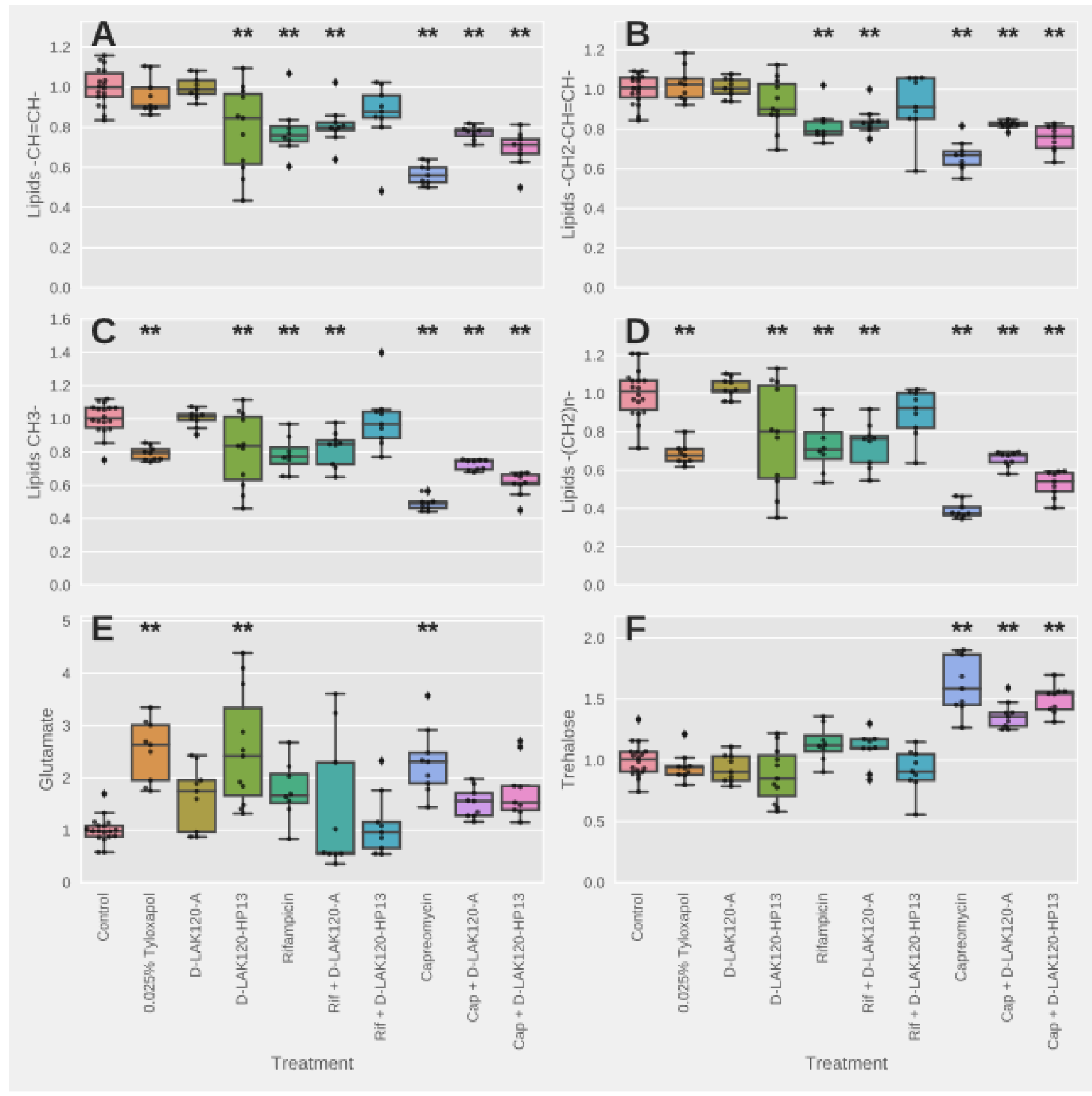
Univariate analysis of relative metabolite levels in *M. smegmatis* mc^2^ 155. Significant differences with respect to unchallenged bacteria, as determined by one-way ANOVA with Tukey post-hoc test, are indicated **. Other significant differences are described in the text.

However, the magnitudes of the responses vary substantially and there are also qualitative differences, most notably since both rifampicin and capreomycin cause reductions in resonances associated with unsaturated alkyl chains. It is notable that the fold changes in the differing lipid resonances are again similar. This is consistent with a reduction in the number of intact mycolic acid molecules rather than a shortening in the length of the mycolic acid alkyl chain. Since resonances for both saturated and unsaturated hydrocarbons are reduced as well as those for the R_2_CH-OH and R_2_CH-COOH protons a general reduction in mycolic acid can be inferred.

Mycolic acids are an important component of the mycomembrane. The inner leaflet is formed of mycolic acid linked to arabinogalactan which is in turn linked to peptidoglycan. The outer leaflet is also formed of lipids based on mycolic acid, but these are considered free to diffuse laterally. These lipids are trehalose dimycolate (TDM) and trehalose monomycolate (TMM). Interestingly, only challenge with capreomycin induces any change in signal intensity attributable to trehalose (Fig. 4F). The significant increase is of a similar magnitude to the decrease observed for the mycolic acid resonances indicating that the ratio of mycolic acid to trehalose is substantially altered in *M. smegmatis* following challenge with capreomycin. This would be consistent with a shift in the balance of TDM and TMM, to favour the latter, in the outer leaflet of the mycomembrane.

The scale of the reduction in the mycolic acid resonances caused by capreomycin (Fig. 4F) is much greater than that caused by rifampicin (Fig. 4E) or even tyloxapol (Fig. 4B). When changes in mycolic acid lipid resonances are compared across the various challenges (Fig. 5A-D) it is clear capreomycin has a substantial impact on those from both saturated and unsaturated hydrocarbons and that while significant (*p* < 0.05), the reductions in intensity due to rifampicin are much more modest. Similarly, tyloxapol causes a modest reduction in resonances from saturated hydrocarbons (Fig. 5C/D) but has no significant effect on those from unsaturated hydrocarbons (Fig. 5A/B). Unlike capreomycin, neither rifampicin nor tyloxapol induce a significant increase in resonances attributable to trehalose (Fig. 5F). Remodelling of the mycolic acid induced by rifampicin and tyloxapol is therefore much subtler than that induced by capreomycin and this is presumably the origin of the nonsignificant changes in membrane physical properties as detected above using fluorescence techniques.

The response of *M. smegmatis* mc^2^ 155 to challenge with ¾ MIC of the two D-LAK peptides is quantitatively and qualitatively different to the response to either rifampicin or capreomycin. Although the OPLS-DA models for challenge with either peptide indicate a significant response when all metabolites are considered (Table 3; Fig. S3), the magnitude of changes in individual metabolites is low and few changes pass a stringent significance threshold (Fig. 4C/D). When comparing individual metabolites across the various conditions, both peptides trigger an increase in glutamate in the bacteria (*p* <0.05), a similar effect is seen for capreomycin but not rifampicin (Fig. 4E). In contrast with capreomycin, neither peptide triggered any change in trehalose (Fig. 5F). The proline containing, and proline free peptides can themselves be distinguished by the absence of any changes in lipid resonances in response to D-LAK120-A while significant (*p* < 0.05) reductions in lipid resonances are observed for both unsaturated and saturated hydrocarbons in response to D-LAK120-HP13 (Fig. 5A-D).

HR-MAS ^1^H NMR data was also obtained for *M. smegmatis* mc^2^ 155 challenged with combinations of each D-LAK peptide and either rifampicin or capreomycin. For synergistic combinations the amount of each antibiotic was substantially lower than that used for each antibiotic alone.

Modest synergism exists between D-LAK120-A and capreomycin but there is indifference or even antagonism between D-LAK120- A and rifampicin. D-LAK120-A mitigates the reduction in saturated and unsaturated hydrocarbons and increase in trehalose due to capreomycin (*p* < 0.05) (Fig. 5A-D, Fig. S1A, Fig. S2A/B) but not the increase in glutamate (Fig 5E). The responses of *M. smegmatis* mc^2^ 155 to rifampicin or rifampicin/D-LAK120-A are indistinguishable (Fig. 5, Fig. S1C, Fig. S2).

D-LAK120-HP13 has a mostly additive effect when used in combination with capreomycin but experiences modest synergism with rifampicin. D-LAK120-HP13 has a weaker impact on metabolic changes induced by capreomycin or rifampicin. Although the magnitude of reduction of lipid resonances in D-LAK120-HP13/capreomycin combinations was consistently lower than that observed when capreomycin was used alone (Fig. 5A-D, Fig. S1B, Fig. S2A/B), the mitigation was in no cases significant. In contrast, although the magnitude of changes induced by rifampicin is less, a significant mitigation (*p* < 0.05) was observed for lipid ‒CH3 (Fig. 5C) while for other metabolites the combination of D-LAK120-HP13 and rifampicin did not induce significant changes with respect to the unchallenged bacteria (Fig. 5A/B/D). Notably the Volcano plot of the individual comparison indicates no metabolites were significantly changed when this combination was used (Fig. S1D) in contrast to rifampicin (Fig. 4F) or indeed D-LAK120-HP13 when used alone (Fig. 4D).

*Increased membrane rigidity is associated with altered composition of the mycomembrane* - to better understand the contributions of the various changes in metabolite concentrations on the physical properties of the mycomembrane following challenge with capreomycin and/or D-LAK peptides or growth in the presence of 0.025% tyloxapol, Spearman correlations between individual metabolites and the fluorescence anisotropy recorded for the same samples were determined using partial least squares regression (Fig. 6, Fig. S6). Significant (*p* < 0.0001) correlations were detected between fluorescence anisotropy and each of the metabolites in which substantial changes resulted from challenge with capreomycin, D-LAK120-HP13 or growth in the presence of 0.025% tyloxapol. The strongest of these correlations were negative correlations with saturated hydrocarbons (Fig. 6A, Fig. 6E) while weaker negative or positive correlations were detected with, respectively, unsaturated hydrocarbons (Fig. 6C/D) or trehalose (Fig. 6B). The strength of these correlations may reflect the impact of each component of the mycomembrane in determining its fluidity or this may reflect the observation that, while all three types of resonance are affected by challenge with capreomycin, D-LAK120-HP13 affects only lipid resonances and tyloxapol affects only saturated hydrocarbons in the shorter α chain. Nevertheless, the increased rigidity of the mycomembrane appears to be a direct result of the observed changes in its composition.

*Rifampicin but not capreomycin induce detectable changes in spent media metabolite composition* - Spent bacterial culture supernatants were also analysed by OPLDS-DA (Fig. S7). Comparison of the models indicates that only those conditions where rifampicin is present was any qualitative change in *M. smegmatis* metabolism detected (Fig. S7A). A binary comparison of spent media for *M. smegmatis* grown without challenge with fresh media reveals the bacteria consume glucose and produce tartrate and 2-hydroxyisobutyrate (Fig. S7B). When challenged with rifampicin there is a slight increase in glucose and citrate consumption but substantially reduced production of lactate, alanine, valine and pyruvate and more modest reductions in production of malate and tartrate (Fig. S7C). These changes were muted when rifampicin was applied in combination with D-LAK120-HP13 where increased consumption of glutamate was also observed (Fig. S7D). In no other conditions were any changes in individual metabolites detected that passed the significance threshold.

## Discussion

The combination of HR-MAS ^1^H NMR metabolomics and fluorescent probes sensitive to changes in membrane fluidity and order reveals modest changes in the mycomembrane of *M. smegmatis* mc^2^ 155 in response to challenge with rifampicin but much more dramatic changes in response to challenge with capreomycin. The D-LAK peptides cannot be distinguished based on any change in the physical properties of the mycomembrane but changes in its composition reflect a distinct mechanism for the proline free and proline containing analogues. The combinations of both D-LAK peptides with capreomycin are synergistic in the presence of tyloxapol but when *M. smegmatis* is cultured in the absence of tyloxapol this is attenuated leaving only modest synergy and only with D-LAK120-A and not D-LAK120-HP13; only the latter is modestly beneficial when combined with rifampicin. These findings may provide a rationale for understanding the properties required for D-LAK peptides and related molecules to potentiate the activity of first and second line treatments against mycobacterial infections and understand whether there is more to the activity of either peptide than enhancing penetration of the mycomembrane.

**Figure 6.**
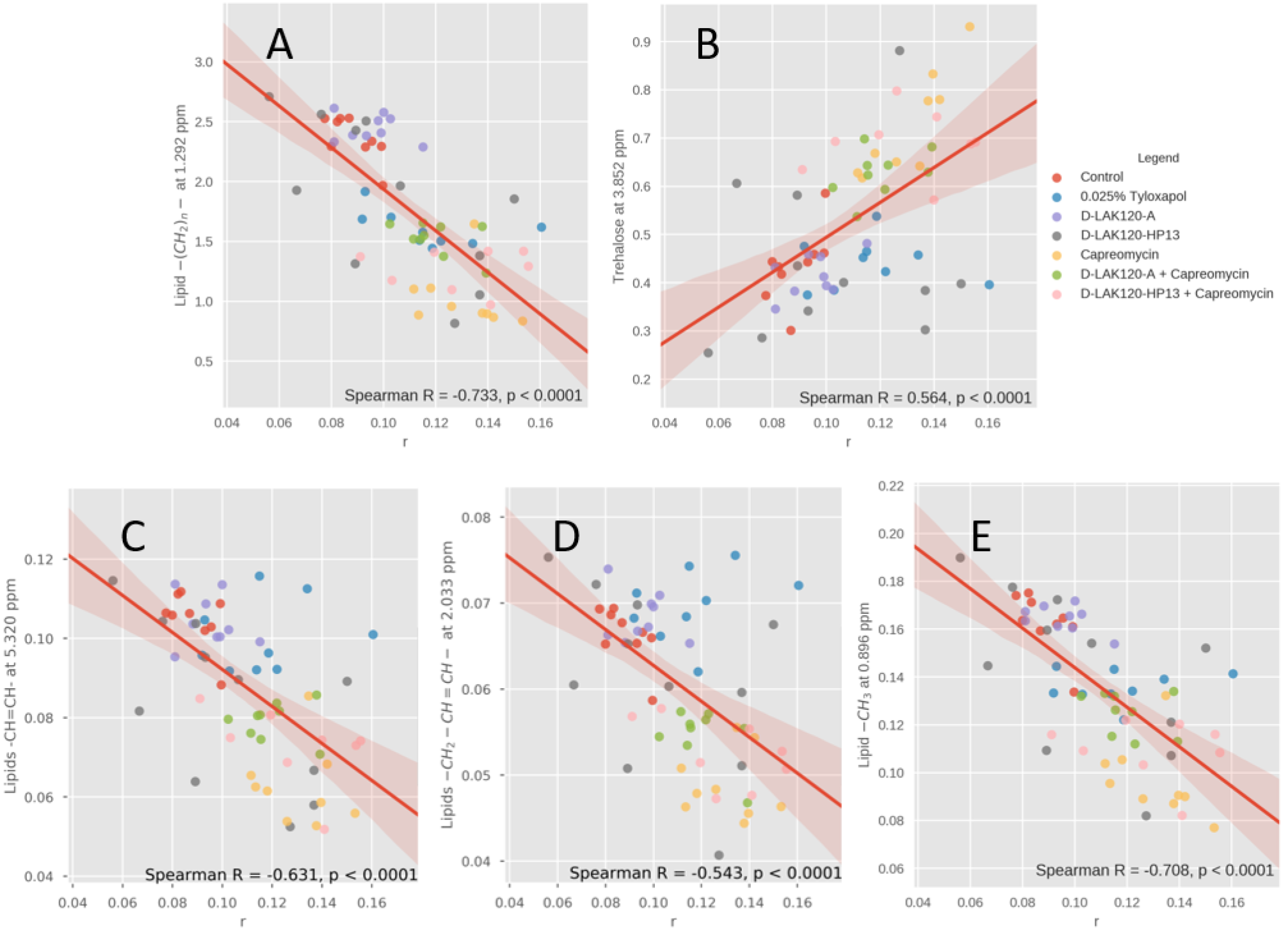
Increased membrane rigidity is associated with altered composition of the mycomembrane. Spearman correlations are shown between DPH anisotropy (r) and ^1^H HR-MAS NMR resonance intensities obtained for *M. smegmatis* mc^2^ 155 grown without challenge or with 0.025% Tyloxapol or ¾ MIC of D-LAK120-A, D-LAK120-HP13, capreomycin, capreomycin + D-LAK120-A or capreomycin + D-LAK120-HP13. Data is shown for lipid -(CH_2_)_n_- (A), trehalose (B), lipid ‒CH=CH- (C), lipid ‒CH2-CH=CH- (D) and lipid ‒CH3 (E).

*Capreomycin acts by inhibiting protein synthesis but induces substantial remodelling of the mycomembrane* - the antibiotic activity of capreomycin against both *M. smegmatis* and *M. tuberculosis* is ascribed to its ability to inhibit protein translation by interfering with the function of ribosomes.^43^ Specifically capreomycin binds across the ribosomal subunit interface using *tlyA*- encoded methylations in both 16S and 23S rRNAs.^44^ A transcriptomic approach to understanding the mechanism of action of capreomycin confirmed the importance of this mechanism but additionally revealed substantial changes in a variety of other gene classes including lipid metabolism, cell wall and cell processes, and intermediary metabolism and respiration in *M. tuberculosis*.^45^ Notably capreomycin was shown to affect the glyoxylate shunt, an alternate pathway to the TCA cycle, with up-regulation of *icl* (Rv0467 - isocitrate lyase), *glcB* (Rv1837c) and *aceAa* (Rv1915) presumably stimulating a process where fatty acids become an important carbon source. Reduced expression of a block of genes associated with the electron transport chain, coding for NADH dehydrogenase or NADH-ubiquinone oxidoreductase, as well as *gdh*, a probable NAD-dependent glutamate dehydrogenase. Though performed in *M. smegmatis* the present study is consistent with these findings; glutamate is seen to accumulate following challenge with capreomycin (and also both D-LAK peptides) while the substantial reductions in mycolic acid is consistent with fatty acids being diverted for use as a carbon source; notably, in contrast with rifampicin challenge, no significant change in individual metabolite levels in spent culture media was detected. Taken together the two studies indicate that there is a substantial shift in metabolism following capreomycin challenge resulting in greater consumption of fatty acids, depletion of mycolic acid and a rigidification of the mycomembrane as trehalose monomycolate predominates over trehalose dimycolate. As such, this response from mycobacteria to capreomycin challenge affords a means of altering the barrier that other antimicrobials may have to cross to reach their targets.

*Rifampicin acts by inhibiting transcription and induces modest remodelling of the mycomembrane* - the antibiotic activity of rifampicin is attributed to its ability to inhibit bacterial DNA-dependent RNA polymerase activity. *M. tuberculosis* is more susceptible to rifampicin then*M. smegmatis* with the latter enjoying a higher level of baseline resistance due to rifampin ADP ribosyltransferase, an enzyme capable of inactivating rifampicin.^46^ However, exposure of rifampicin sensitive *M. tuberculosis* H37Rv to rifampicin induces changes in expression of genes in the same functional classes affected by capreomycin challenge i.e. lipid metabolism, cell-wall and cell processes, and intermediary metabolism and respiration.^47^ Further, rifampicin, and isoniazid and streptomycin, trigger activation of isocitrate lyases in *M. tuberculosis*^48^ suggesting the glyoxylate shunt is a general means of overcoming oxidative stress.^49^ Antimicrobial peptides including LL-37 and pleurocidin have been shown to induce oxidative stress respectively in *Escherichia coli* and *Candida albicans*.^50,51^The modest changes in mycolic acid induced by challenge with rifampicin and D-LAK120-HP13 may therefore be indicative of oxidative stress and reflect penetration of the bacterium; notably D-LAK120-A does not have this effect. The response of *M. smegmatis* to rifampicin differs from that to capreomycin quantitatively in the HR-MAS study but also qualitatively in the study of spent culture composition where evidence for an alternate strategy to overcoming oxidative stress emerges. This would be consistent with *M. smegmatis* altering its metabolism in response to rifampicin in a similar way to that used in response to capreomycin but being less reliant on metabolism of fatty acids.

*Distinct membrane activity underpins synergistic combinations* - for antibiotics to have additive or synergistic effects when used in combination they need to avoid both interactions that reduce the amount of each agent reaching their target(s) and triggering responses in the bacteria that counter the activity of the other agent. For combinations of antibiotics to have synergistic effects, additionally they need to act on different targets or have distinct effects on the same target such that the effect of each individual antibiotic is enhanced. The HR-MAS ^1^H NMR approach enables investigation of whether antibiotics do indeed differ in their effect on the mycomembrane. This is achieved by obtaining a bacterial perspective of the challenge and inferring differences in activity of each antibiotic from differences in the response of the bacteria to each challenge. Each of capreomycin, rifampicin, the D-LAK peptides and tyloxapol induce qualitatively and/or quantitatively different responses from *M. smegmatis* but there is nevertheless considerable overlap.

Tyloxapol improves the potency of rifampicin and D-LAK peptides when used alone but the combination of D-LAK peptides and rifampicin is indifferent or even antagonistic in the presence of tyloxapol. Since both rifampicin and tyloxapol induce membrane remodelling in *M. smegmatis* one might speculate that rifampicin and tyloxapol combine to produce substantial change in membrane that inhibits the ability of D-LAK peptides to disrupt or penetrate membrane. In the absence of tyloxapol, the change in membrane properties induced by rifampicin are more modest and this may explain why combinations of rifampicin with D-LAK120-A fare worse than those containing D-LAK120-HP13 if the proline modification enables this analogue to better penetrate the mycomembrane.

Both D-LAK peptides interact synergistically with capreomycin, albeit modestly. The interaction is enhanced however when *M. smegmatis* is cultured in the presence of 0.025% tyloxapol. While tyloxapol enhances the action of both peptides and rifampicin, the MIC for capreomycin is not reduced and may even be higher. It is unlikely therefore that tyloxapol improves access for capreomycin though it may perform this function for either D-LAK peptide. Although the difference between the two peptides in terms of their interaction with capreomycin is small, D-LAK120-A consistently outperforms D-LAK120-HP13. Both peptides seemingly have a similar effect on capreomycin activity but the poorer synergy between D-LAK120-HP13 and capreomycin may be related to the ability of both agents to trigger changes in the mycomembrane. Since both capreomycin and rifampicin trigger changes in the mycomembrane composition, albeit with very different magnitudes of response, this argument seemingly runs counter to the idea that membrane interactions will determine the ability of such molecules to interact synergistically against *M. smegmatis*. The answer to this may lie in the distinct physical properties of capreomycin and rifampicin. Capreomycin is a cationic molecule with a physiological net positive charge of four. Rifampicin is also cationic but its physiological charge is only one. Both molecules must penetrate the bacteria to exert their main antibiotic effects and hence the D-LAK peptides, with a net positive charge of nine and a higher affinity for anionic components of the mycomembrane, may provide an important screen for interactions between the membrane and either capreomycin or rifampicin. Capreomycin, with the higher charge, would benefit more from this screen and this may underlie the observed synergy rather than any induced change in mycomembrane biophysical properties.

## Conclusion

The two D-LAK peptide analogues can be distinguished based on the different response of *M. smegmatis* to AMP challenge and their different abilities to enhance the activity of rifampicin and capreomycin; two anti-mycobacteria drugs with distinct mechanisms of action but related in their requirements for efficacy and the bacterial response to their activity. In addition to distinguishing anti-mycobacterial drugs the combination of HR-MAS ^1^H NMR and fluorescence spectroscopy reveals how bacterial responses to each agent; remodelling of the mycomembrane may affect their ability to act in combination. In addition to consideration of the mode of action and physical properties of each antibiotic, understanding such effects may be essential in designing effective antibiotic combinations.

## ASSOCIATED CONTENT

**Supporting Information.** Supplementary figures are available as described in the text.

## Author Contributions

AJM and DM wrote the main manuscript text and prepared all figures. DM, AJM, BDR and JKWL designed experiments. BDR and JKWL provided materials. DM and TK performed and/or analysed NMR metabolomic experiments/data. DM conducted susceptibility testing and obtained electron microscopy and fluorescence microscopy images. All authors approved the manuscript.

### Notes

The authors declare no competing financial interest.

## ACKNOWLEDGMENT

This work was supported by a joint HKU-KCL studentship for DM awarded to JKWL, the Wellcome Trust (Capital Award for the KCL Centre for Biomolecular Spectroscopy). This work was supported by the Francis Crick Institute through provision of access to the MRC Biomedical NMR Centre. The Francis Crick Institute receives its core funding from Cancer Research UK (FC001029), the UK Medical Research Council (FC001029), and the Wellcome Trust (FC001029). We are grateful to Drs Tom Frenkiel, Alain Oregioni and Andrew Atkinson for their assistance with NMR instrumentation. We also acknowledge the assistance of the Li Ka Shing Faculty of Medicine Faculty Core Facility, and the Electron Microscope Unit at the University of Hong Kong.

